# Practical Identifiability in the Frame of Nonlinear Mixed Effects Models: the Example of the *in vitro* Erythropoiesis

**DOI:** 10.1101/2021.03.01.433388

**Authors:** Ronan Duchesne, Anissa Guillemin, Olivier Gandrillon, Fabien Crauste

## Abstract

Nonlinear mixed effects models provide a way to mathematically describe experimental data involving a lot of inter-individual heterogeneity. In order to assess their practical identifiability and estimate confidence intervals for their parameters, most mixed effects modelling programs use the Fisher Information Matrix. However, in complex nonlinear models, this approach can mask practical unidentifiabilities. Herein we rather propose a multistart approach, and use it to simplify our model by reducing the number of its parameters, in order to make it identifiable. Our model describes several cell populations involved in the *in vitro* differentiation of chicken erythroid progenitors grown in the same environment. Inter-individual variability observed in cell population counts is explained by variations of the differentiation and proliferation rates between replicates of the experiment. Alternatively, we test a model with varying initial condition. We conclude by relating experimental variability to precise and identifiable variations between the replicates of the experiment of some model parameters.

## 1 Introduction

Inter-individual variability is ubiquitous in biology, from the fluctuations of molecular contents across populations of single cells [1], to the variations of physiologial parameters between whole organisms [2]. This variability has uncountable consequences, for instance at the scale of developmental [3], ecological or evolutionary processes [4, 5].

As a result, one often faces significant amounts of variations between replicates of the same biological experiment, which we will refer to as *experimental variability*. This variability can be taken into account by deterministic dynamical models of the biological system, as a random variation around its predicted behaviour [6, 7]. Such models thus disregard the fact that variability is inherent to the biological nature of the system under study.

Another difficulty that can arise from this approach is when some parameters of the model are unidentifiable. A parameter is said identifiable when a particular measurement of the model output (potentially affected by measurement error) is associated to a unique parameter value [6]. Otherwise it is unidentifiable. The model itself is said identifiable if all its parameters are identifiable, and unidentifiable if at least one of its parameters is unidentifiable.

More precisely, a model can be unidentifiable for several reasons. If some parameters are redundant, meaning that they can be varied together in such a manner that the model output is kept constant, they are called structurally unidentifiable. If the data quantity (sample size) or quality (measurement error) are insufficient to precisely infer some parameter values, these parameters are said practically unidentifiable. It should be noted that while all practically identifiable parameters of a model are also structurally identifiable, the converse is not necessarily true (see for instance the recent review on identifiabilty by Wieland *et al*. [8]). For this reason, the focus of this paper is on practical identifiability, and unless stated otherwise we will be referring to the practical identifiability of model parameters.

When the parameters of a model have a precise physical or biological interpretation, it can be tempting to use their estimates to formulate predictions about the system. However, as these estimates are not uniquely determined in unidentifiable models, an unidentifiable model should never be used for predictive purposes [9].

In order to interpret experimental heterogeneity, we propose to use nonlinear Mixed Effects Models (MEM). In particular, we are interested in the identifiability of such models. MEM work by applying the same mathematical model to all the individuals of the population, with different parameter values for each individual, and thus have been used in a variety of fields involving inter-individual variability [10]. This approach allows to assign different levels of variability for each parameter by making the distinction between *population parameters*, that are the means and variances of the parameter values across the whole population, and *individual parameters*, that are the precise parameter values assigned to each individual.

In the context of experimental variability, one might for instance consider all replicates of the experiment as individuals coming from the same population (the theoretical population of all the possible outcomes of the experiment). Assigning different parameter values in a dynamic model for each individual (*i*.*e*. each replicate of the experiment) would then allow to assess which parameters of the model are mostly affected by experimental variability (*i*.*e*. which parameters are the most variable between individuals). The question that naturally arises is whether or not such a model would be identifiable.

In general, one argument in favour of the use of MEM is the fact that using the population distribution as a prior can help with the estimation of the individual parameters, thus improving their practical identifiability [11]. This rationale is based upon the fact that most MEM calibration methods estimate the population parameters in a first step, using the data from the whole population, and then estimate individual values for every parameter in a bayesian way, an approach referred to as *Empirical Bayesian Estimation* [12].

However, this first intuitive argument on parameter identifiability in MEM is somewhat challenged by another consideration: the intricate definition and estimation of population and individual parameters might in fact complicate the assessment, and even the definition, of parameter identifiability in MEM [13]. As a consequence, the identifiability of MEM parameters is of critical importance to their widespread applications, and should not be neglected [9, 14]. In general, practical parameter identifiability depends on the precise definition of the model parameters (in the case of MEM, that is the definition of the distributions of the individual parameters across the population), together with the quantity and quality of the experimental data used for calibration [6]. Several kinds of approaches for structural identifiability analysis, that were developped for models without mixed effects, have been adapted in a mixed effect context [15, 16]. Regarding practical identifiability analysis, two different kinds of empirical approaches are reported for MEM [13], though with a lot of potential refinements.

First, the *Fisher Information Matrix* (FIM), which is computed from the Hessian of the likelihood, estimated at the optimal parameter set, allows for a quadratic approximation of the likelihood surface near its optimum. This in turn, allows to infer confidence intervals for any parameter at any level, provided that the FIM is non-singular [17, 18]. In this setting, a singular FIM indicates that some parameters are structurally unidentifiable. Conversely, a near-singular FIM could indicate that some parameters are practically unidentifiable. However, the quadratic approximation of the likelihood surface might mask practically unidentifiable parameters in the case of nonlinear, partially observed models. In extreme cases, the FIM can even make some practically unidentifiable parameters appear as identifiable [6]. As a consequence, while the FIM is still relevant to assess the structural identifiability of models, it is completely inadequate to study practical identifiability. Since our focus is on the practical identifiability of MEM, we will not use the FIM to assess the identifiability of our models. Other methods have been developed for models without mixed effects, such as the profile likelihood [6], but to our knowledge they have not yet been implemented in any of the existing software for MEM calibration. Given the widespread use of these software for the calibration of MEM in pharmacology and personalized medicine [19], it would be particularly interesting to be able to empirically study the practical identifiability of MEM, directly from the calibration software.

Secondly, one might run the estimation algorithm several times, using a sample of initial guesses for the parameter values [13]. In that case, the convergence of the algorithm to a unique likelihood optimum, with different optimal parameter values, indicates that the parameters are unidentifiable. We refer to this approach as *Initial Guess Sampling* (IGS). It has also been termed the *multistart approach*, and the samples of estimated parameter values that it provides do not contain any information regarding the variance of the estimation or the confidence intervals of the parameters [20]. Since this method requires a potentially large sample of runs of the estimation algorithm, it is more costly in terms of computational power than a simple evaluation of the FIM eigenvalues. As a consequence, most approaches to the identifiability analysis of MEM rely on the FIM [10, 13, 14, 21–23], despite its proven inaccuracy at assessing practical identifiability [6].

Since the practical identifiability of a model depends on both the definition of the model and the data used to calibrate it, there are two broad classes of approaches for dealing with an unidentifiable model. On one hand, it is possible to increase the amount of data available for parameter estimation. For instance measuring an additional, previously unobserved quantity might remove structural unidentifiabilities. It is also possible to use the tools of experimental design [22, 23] to define a new set of more informative experiments, that can then be used to estimate new parameter values and assess their identifiability. However, experimental design approaches are approximate and there is no *a priori* proof that performing the optimal experiment will actually make the model identifiable [24, 25]. On the other hand, it is also possible to change the definition of the model parameters, in order to simplify the estimation for the other parameters [26, Section 10.2], an approach which we refer to as *model reduction*. For instance, reparameterizing the model in terms of the estimable parameter combinations would remove any structural unidentifiabilities, while potentially sacrificing the biological interpretation of these parameters. It is also possible to constrain the value of some unidentifiable parameters (for instance setting them to zero) in order to simplify the estimation task. But then, which criterion would allow us to choose which parameters to remove from the model?

This consideration is particularly important for MEM, as all individual parameters might not have the same variance across the population. Thus, the size of the sampled dataset (in terms of the number of individuals) is critical for the estimation of the population variances, since a sampling bias in a small dataset could mask the variance on some parameters. From the point of view of experimental design, the determination of the necessary sample size in order to guarantee a certain level of confidence on all parameters in MEM is a central question, which has already been covered to a certain extent [22, 27], using geometric features of the likelihood surface approximated from the FIM [28, section 10.5.3]. From the point of view of model reduction, it would be tempting to remove the unidentifiable variances by setting them to zero, potentially improving the estimation of the other parameters of the MEM without affecting the quality of the model fit to the data. In some cases however, adding a random effect or a covariate to a MEM could improve parameter identifiability [26, Section 5.1], as it might split out combinations of structurally unidentifiable parameters.

In this paper, we adress these questions using a MEM of the *in vitro* erythropoeisis that we adapt from a previous model, proven to relevantly reproduce the dynamics of single replicates of the experiment [29]. This MEM accounts for experimental variability by assigning different parameter values for proliferation and differentiation in each replicate of an identical experiment. We assess its identifiability using a multistart approach, based on extensive parameter estimations with the MEM calibration software Monolix [30]. Then, we reduce the model in order to make it identifiable, using the correlations between the estimated parameter values. Alternatively, we test whether or not the observed variations in the outcome of our experiment could be explained by variations in the initial condition of the experiment rather than variations of the differentiation and proliferation dynamics. Our final model associates different levels of variability for each dynamic parameter, which allows us to identify which features of the erythroid differentiation are the most variable from experiment to experiment. Moreover, this work proposes a multistart approach for MEM identifiability analysis, which appears as a promising alternative to the FIM.

## 2 Materials and methods

### T2EC cell culture

The experimental setting from which all the data used in this study were obtained consists in a culture of 25000 chicken erythroid progenitors called T2EC that were extracted from the bone marrow of 19 days-old embryos of SPAFAS white leghorn chickens (INRA, Tours, France). They may either be maintained in a proliferative state or induced to differentiate into mature erythrocytes depending on the medium in which they are grown, as previously described [29, 31–33].

In the self-renewal medium (referred to as the LM1 medium) the progenitors self-renew, and undergo successive rounds of division. Its composition is given in Table S1 in the Supplementary Materials. Cell population growth was evaluated by counting living cells in a 30 µL sample of the 1mL culture using a Malassez cell and Trypan blue staining (SIGMA), which specifically dyes dead cells, each 24h after the initial sowing of 25000 cells in the culture, as previously described [29, 31–33].

T2EC can be induced to differentiate by removing the LM1 medium and placing cells into 1mL of the differentiation medium, referred to as DM17. Its composition is given in Table S1 in the Supplementary Materials. Upon the switching of culture medium, a fraction of the progenitors undergoes differentiation and becomes erythrocytes. The culture thus becomes a mixture of differentiated and undifferentiated cells, with some keeping proliferating. Cell population differentiation was evaluated by counting differentiated cells in a 30 µL sample of the culture using a counting cell and benzidine (SIGMA) staining which stains haemoglobin in blue. A parallel staining with trypan blue still gives access to the overall numbers of living cells, as previously described [29, 31–33].

Consequently, the data available from this experiment are the absolute numbers of differentiated cells, as well as the total number of living cells (which comprises both self-renewing and differentiated cells) each 24h after the initial sowing of 25000 cells in the culture. The data presented on Figure 1 are the total number of living cells in the culture, and the fraction of differentiated cells in 7 independent replicates of the experiment.

**Figure 1:**
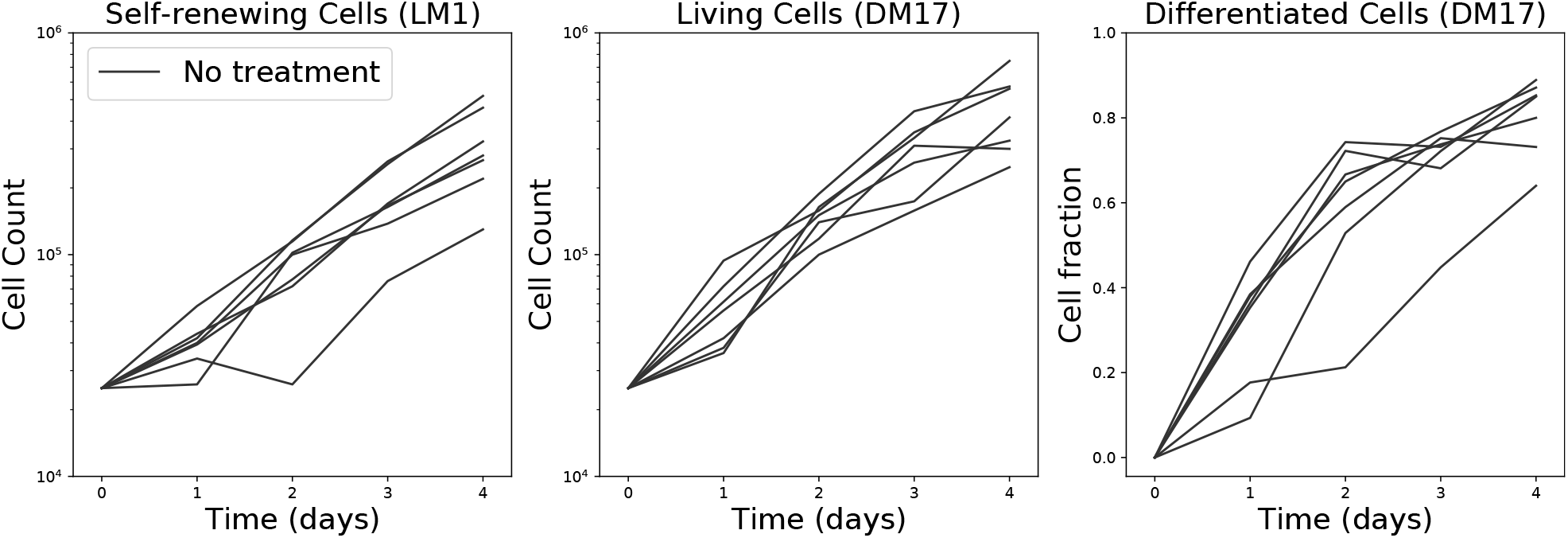
Data used in this study. They comprise the total number of living cells in the LM1 and in the DM17 media (in log-scale), as well as the number of differentiated cells in DM17 (represented as a fraction of the total number of cells) in 7 independent replicates of the experiment.

### 2.1 Modelling framework

A Mixed Effects Model (MEM) is defined as the combination of three components. The structural model describes the dynamic process at play in each individual. The parameter model, or individual model, describes how the parameters of the structural model vary from individual to individual. Finally, the observation model, or error model, describes how the predicted outcome of the model for each individual differs from the observation.

#### 2.1.1 Dynamic model

The SCB model, that we previously described [29], faithfully reproduces the dynamics of T2EC proliferation and differentiation by accounting for 3 cellular states (Figure 2). The self-renewing state S describes the state of cells in the LM1 medium, where they can only proliferate or die. The differentiated state B (which stands for *Benzidine-positive*) describes mature erythrocytes in the DM17 medium. Lastly, in the committed state C, cells have not finished differentiating, but cannot go back to self-renewal anymore, so that they are committed to differentiation. The dynamics of these three compartments are given by the equations:

**Figure 2:**
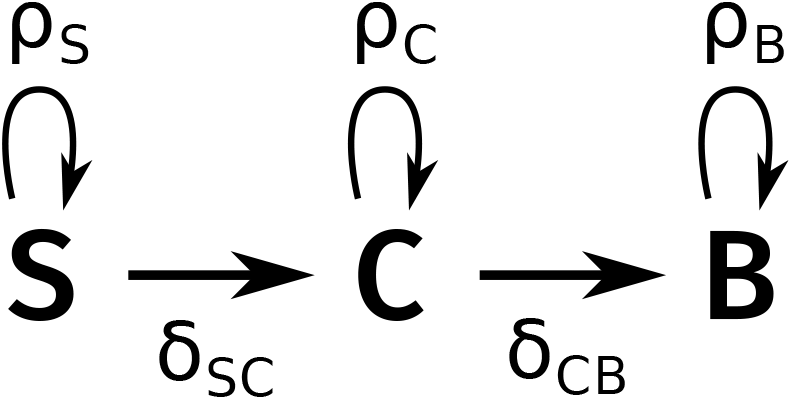
Diagram of the dynamic model. S: self-renewing cells. C: committed cells. B: Benzidine-positive (*i*.*e*. differentiated) cells. *ρ*_*i*_ denotes the proliferation rate of compartment *i* and *δ*_*ij*_ is the differentiation rate of compartment *i* into compartment *j*.

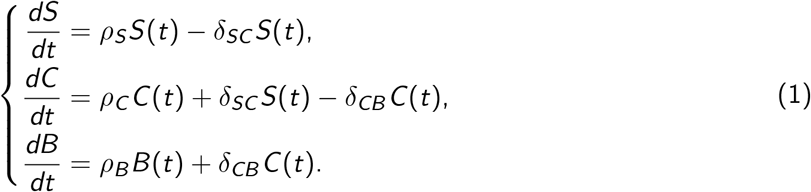

It is characterized by the set (*ρ*_*S*_, *δ*_*SC*_, *ρ*_*C*_, *δ*_*CB*_, *ρ*_*B*_) of five dynamic (or kinetic) parameters, where *ρ*_*i*_ is the net growth rate of compartment *i*, which might be positive or negative depending on the net balance between cell proliferation and cell death, and *δ*_*ij*_ is the differentiation rate of compartment *i* into compartment *j*, which must be positive.

Moreover, it should be noted that differential system (1) is fully linear, and that its matrix is lower-triangular, which makes it easily solvable analytically. Its simulation is thus very fast. The detail of the analytical solutions to this system is given as supplementary material in [29].

Finally, not all variables in the models can be measured through the experiments that we presented in Section 2, and we only have access to three observables of the system: the number of living cells in LM1 (which we denote as *S* since there are only self-renewing cells in LM1), the number *T* of living cells in DM17, and the number *B* of differentiated cells in DM17 (it is null in LM1). Unless stated otherwise, we will always consider that the initial condition is fixed by the experimentalist so that the initial state of the observables is: (*S*_0_, *T*_0_, *B*_0_) = (25000, 25000, 0).

#### 2.1.2 Parameter model

In order to describe inter-individual variability in the parameter values of the SCB model, we consider at first that the five kinetic parameters can vary between every individual. The two differentiation rates *δ*_*SC*_ and *δ*_*CB*_ must be positive, and the net proliferation rates *ρ*_*S*_, *ρ*_*C*_ and *ρ*_*B*_ can be positive or negative. In order to respect these bounds on the individual parameter values, we use a combination of normal and lognormal distributions across the population:

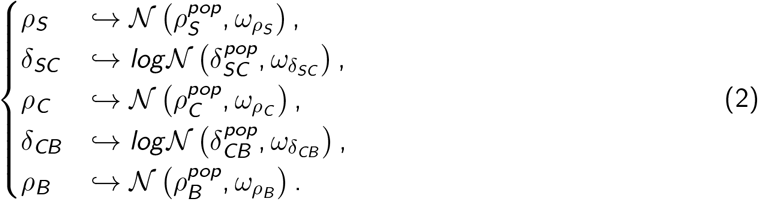

This parameter model has 5 fixed effects (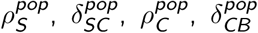,and 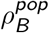), which quantify the average behaviour of the population, and 5 random effects (associated with the standard deviations 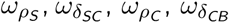, and 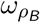).

#### 2.1.3 Error model

In order to account for experimental errors in the measurement of the observables, MEM include an error model, or observational model, which describes the statistical fluctuation of the model prediction around the observation. We previously demonstrated that the proportional error model is the best to describe the prediction error of the SCB model [29]. It is defined by:

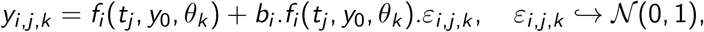

where *y*_*i,j,k*_ marks the measurement of the *i* ^*th*^ observable, at the *j*^*th*^ timepoint, on the *k*^*th*^ individual, and *f*_*i*_ marks the model prediction for the *i* ^*th*^ observable from System (1), which depends on time *t*, the initial condition *y*_0_, and the individual parameters *θ*_*k*_. Finally, *b*_*i*_ denotes the proportional error parameter for the *i* ^*th*^ observable, which quantifies the standard deviation of the prediction error, and *ε*_*i,j,k*_ is the individual weighted residual of the model for individual *k*, at time *t*_*j*_, for observable *i*. The proportional error model introduces one additional parameter *b*_*i*_ for each observable, resulting in three error parameters for our SCB model.

Together with the dynamic model of System (1) and the parameter model of Equation (2), this error model defines our first version of a MEM for the *in vitro* erythropoiesis. Since all other MEM in this manuscript will have the same dynamic and error components, we will omit them from now on, and will define each MEM by its parameter equation only, such as Equation (2).

### 2.2 Parameter estimation

We used the Stochastic Approximation version of the Expectation-Maximization (SAEM) algorithm [34] implemented in Monolix [30] to estimate the parameters of our MEM (Table S2). To avoid potential local likelihood optima and ensure the convergence of the algorithm to the global optimum, we performed the estimation 50 times using the Monolix Convergence Assessment tool, with independent uniformly sampled initial guesses for the parameter values (Table S3). For the fixed effects, we sample the initial guess in an arbitrarily large interval (Table S3). Thus most initial guesses will be wrong, and potentially far from the true value. To ensure convergence in these conditions, we set a high initial variance and error parameter values (Table S3). This ensures that the first step of SAEM can sample individuals across all the parameter space, which will allow for a subsequent improvement of the estimates both for the population averages and variances. This is a multistart approach that we refer to as *Initial Guess Sampling*.

### 2.3 Model selection

In order to select which SAEM runs converged to the global optimum, we used Akaike’s weights [35]:

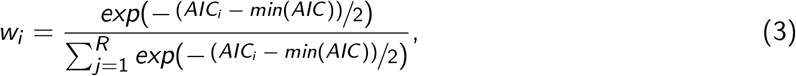

where *w*_*i*_ is the Akaike’s weight of the i-th run, *AIC*_*i*_ is its Akaike’s Information Criterion and *R* is the number of competing models. The Akaike’s weight of a given model in a given set of models can be seen as the probability that it is the best one among the set [35]. In this setting, selecting the best models of a set of models means computing their Akaike’s weights, sorting them, and keeping only the models whose weights add up to a significance probability (in this manuscript, 95%).

In MEM, one might either choose to use the marginal or the conditional AIC depending on the context [36]. They differ by the corrective term that they introduce in the likelihood. However, we will essentially use the AIC and the corresponding Akaike’s weights for selecting models with the same structure and different likelihoods, such as the 50 runs of SAEM that we perform on the same model to assess its convergence. For this reason, the choice of mAIC or cAIC is not relevant to our study, and we will use the marginal 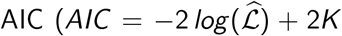, where 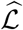 is the maximum likelihood and *K* is the number of population parameters), that is computed by Monolix by default [10].

In order to select models with different structures, *i*.*e*. models that would differ by the definition of their fixed or random effects, we use the BIC that has been derived for MEM [37, 38].

### 2.4 Identifiability analysis

#### 2.4.1 Population parameters

We use an approach based on repeated parameter estimations, starting from different initial guesses, to empirically assess the identifiability of our MEM. In this case, convergence to different parameter values with the same likelihood indicates unidentifiability [13]. This approach has also been termed the *multistart approach*, for instance by [20]. We will refer to the multistart approach as *Initial Guess Sampling* (IGS) in this manuscript.

Our approach to IGS is the following:

1. We perform a random sampling of the initial parameter guesses and run SAEM for each of the initial guesses, using the Monolix Convergence Assessment tool. This provides us with a sample of optimal parameter values.
2. We test the convergence of the SAEM runs: we only want to consider the runs which reached the global optimum. To this end, we use a selection criterion (*w*_*AIC*_) to keep only the runs that converged to the lowest likelihood values.
3. We compare the parameter values of these convergent runs. If they are different, then the model is unidentifiable, as several different parameter values give the same likelihood.

It should be noted that since multistart approaches do not provide any information on the parameters estimation error or confidence intervals [20], this approach do not allow for any statistical testing of parameter identifiability. The distribution of estimated values can rather be used to design a diagnostic plot of population parameter identifiability, showing which parameters vary the most between the convergent runs, and are thus the most poorly estimated. We propose to visualize the distributions of estimated values as a boxplot normalized by their median in order to display this information.

#### 2.4.2 Individual parameters

Individual parameters are estimated using an empirical bayesian approach, where the population distribution of the parameters serves as a prior balanced by the individual data. In the case of unidentifiable individual parameters, the experimental data do not provide enough information to determine them precisely. Then, the posterior distribution is very close to the prior, resulting in individual parameters being estimated as their population mean.

More precisely, this principle that the posterior matches the prior for unidentifiable parameters holds under two conditions [39]: first, the prior distributions must be independent, second, the parameter space must be a product space. When one of these two conditions is not met, it is possible that the posterior distribution will differ significantly from the prior even for unidentifiable parameters [39].

In the case of our model, the prior distributions of the individual parameters, which are the population distributions defined in System (2), are independent, since the variance-covariance matrix of the random effects is diagonal. Moreover, each individual parameter is either real (net self-renewal rates) or positive (differentiation rates), and thus the individual parameter space is a product space, namely 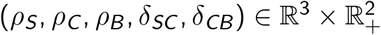

Consequently, we can assess the identifiability of the individual parameters by measuring the overlap between the prior and posterior distributions. This phenomenon is summarized by a scalar criterion called the *η-shrinkage* [12, 40]:

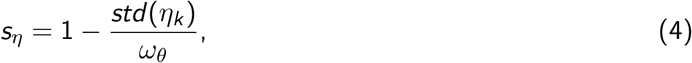

where *std* (*η*_*k*_) is the standard deviation of the estimated individual random effects in the population, and *ω*_*θ*_ is their theoretical standard deviation. In the case where information about a parameter is insufficient, the random effects on this parameter shrink toward 0 in the population, and thus *s*_*η*_ increases. Equation (4) also implies that shrinkage values vary from parameter to parameter, and that some parameter might be more poorly characterized than the others.

Simulation studies have shown that shrinkage has a variety of effects on the model diagnostics, starting from 30% shrinkage [12]. High shrinkage values affect the correlations between random effects and covariates, as well as the correlations between the random effects themselves. It can also affect the detection of structural model specification.

In this paper, we will use the 30% limit introduced in [12] as a rule of thumb to consider that the individual parameters are well estimated.

## 3 Results

### 3.1 The model is unidentifiable

We estimated the parameter values of Model (2) by using our multistart approach. The distribution of estimated likelihood values over the 50 runs of SAEM is displayed on Figure 3A, showing small variations between the estimated log-likelihood values. Among these 50 runs of SAEM, the 45 associated to the 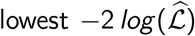 add up to 95 % of Akaike’s weights (Figure 3B). We thus focus on the outcome of these 45 runs in the following.

**Figure 3:**
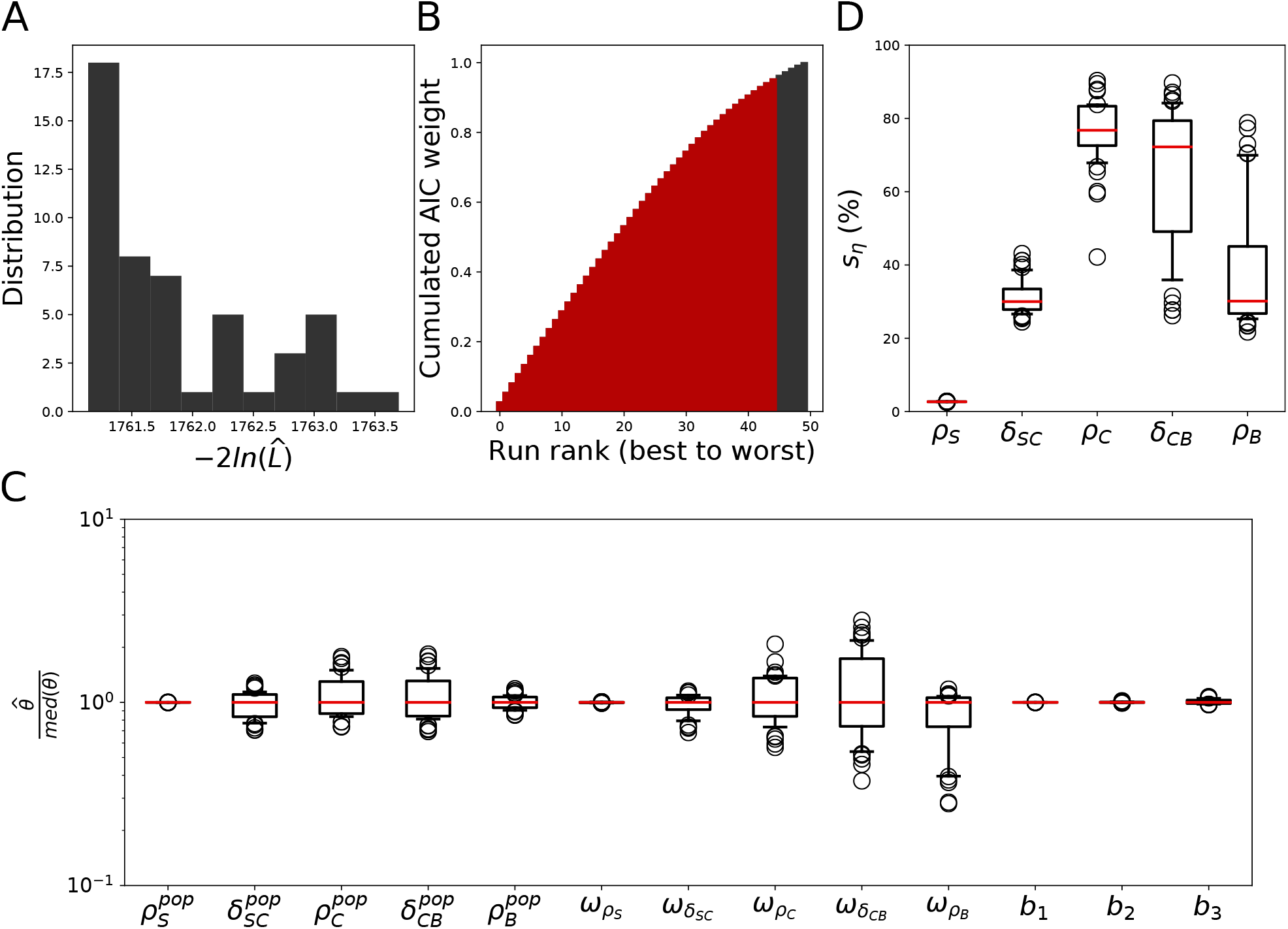
Model (2) is unidentifiable. A: Likelihood distribution over 50 SAEM runs on the Model (2). B: Cumulated AIC weights over the 50 runs of SAEM. The 45 runs associated to the lowest likelihood values (*i*.*e*. those that add up to 95% of the total weight of the 50 runs) are coloured in red. C: Normalized parameter values in the 45 convergents runs of SAEM. Displayed are the distributions of estimated parameter values, normalized by their median. D: Distribution of the *η*-shrinkage values for the individual parameters in the 45 convergent runs of SAEM.

The distributions of estimated population parameter values are represented in Figure 3C. The fixed effect 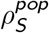, the corresponding variance 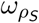 and the three error parameters *b*_1_, *b*_2_ and *b*_3_ are estimated with the smallest variance. For any of the other 8 parameters, the estimated values display more or less variability. For these parameters, the estimation is less reliable. Thus, we conclude that the fixed effects 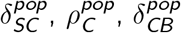 and 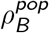 are unidentifiable, as well as the corresponding variances (a total of 8 unidentifiable population parameters).

The shrinkage of the individual random effects is displayed on Figure 3D. The values of shrinkage for *δ*_*SC*_ *ρ*_*C*_, *δ*_*CB*_ and *ρ*_*B*_ range from 20 to 90% depending on the run, which indicates a clear discordance between the population distribution of the parameters and the actual distribution of the individual parameters. We thus conclude that the individual data are not informative enough to estimate all random effects for each individual.

As a consequence, it appears that Model (2) is unidentifiable at the population level as well as at the individual level.

### 3.2 A reduction approach for MEM

#### 3.2.1 Fixed effects: parameter correlations

Figure 4A displays the value of Spearman’s *ρ*^2^, which measures the nonlinear correlation between two variables, for each pair of the 8 unidentifiable population parameters of Model (2). There is a high correlation between 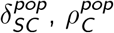 and 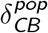 across the runs, which is represented in Figure 4B-C.

**Figure 4:**
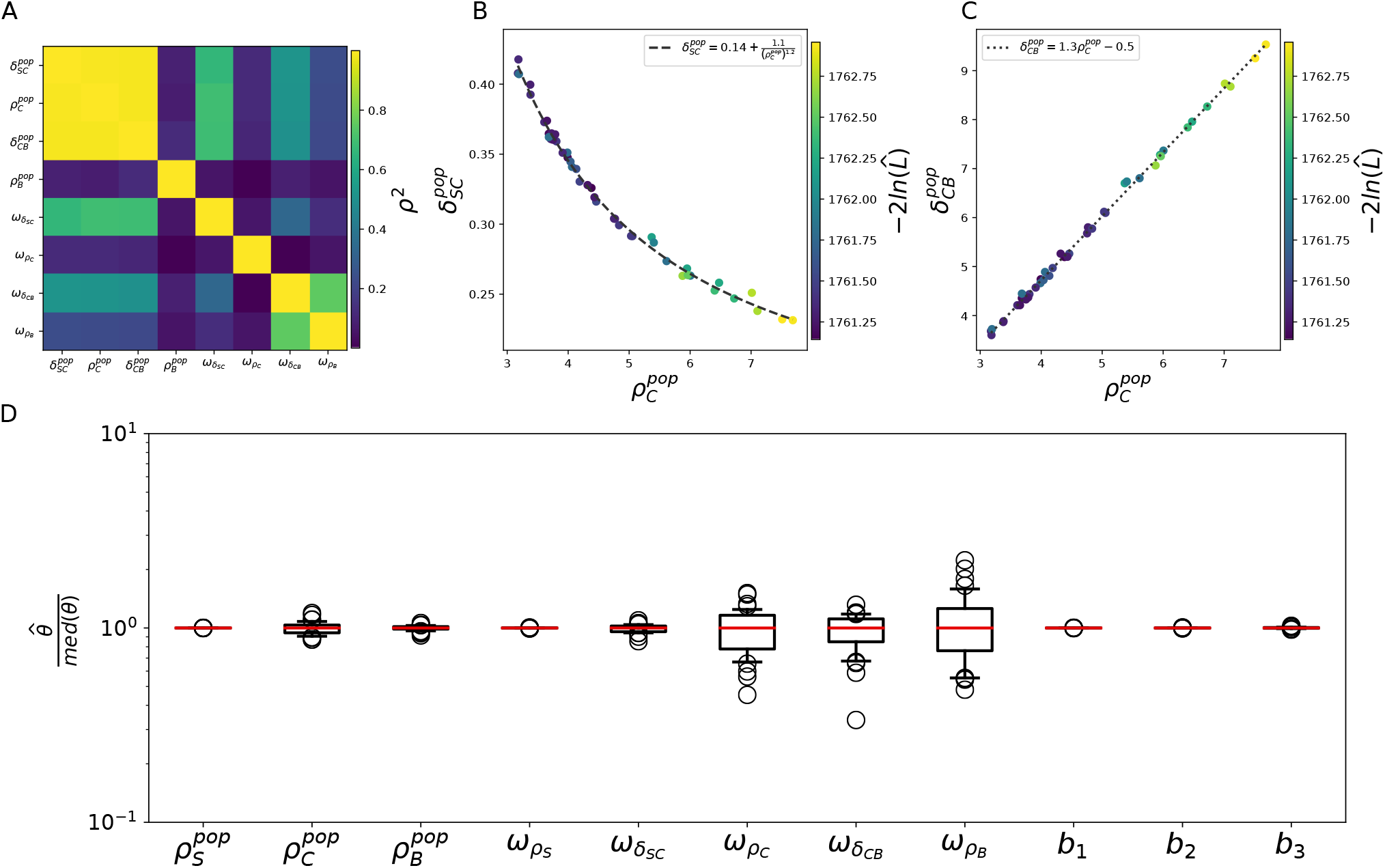
Correlations between the population parameters in Model (2) allow for a reduction of its fixed effects. A: Correlation heatmap (Spearman’s *ρ*^2^) of the 8 unidentifiable population parameters in the 50 runs of SAEM for the initial model. For each pair of unidentifiable population parameters, the heatmap displays the color-coded value of *ρ*^2^. B: Nonlinear correlation between 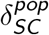 and 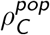. C: Linear correlation between 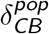 and 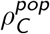. (B-C: Displayed are the estimated population parameter values over the 50 runs of SAEM, color-coded by likelihood.) D: Estimated parameter values in the 36 convergent runs of SAEM for Model (7), with reduced 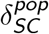 and 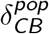. Displayed are the distributions of estimated parameter values, normalized by their median.

These results show that the optimal values of 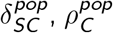 and 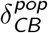 are strongly correlated in the range of values of Figure 4. This range corresponds to the range of estimated values in the 50 SAEM runs. The correlations suggest that if we would replace two of these parameters by their expression as a function of the third one, we would also reduce the number of population parameters to estimate, and still allow them to reach their optimal value.

We found the following expressions for the correlations:

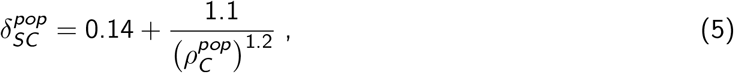

and

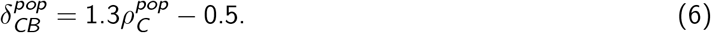

We thus conclude that if we replace 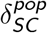 and 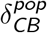 by their expression as a function of 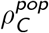 in Model (2), it might help the estimation. Yet, such a reduction might affect the convergence of SAEM because the correlation might not hold outside of the parameter range of Figure 4.

Replacing 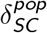 and 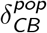 in Model (2) by their expression as a function of 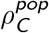, we obtain the following reduced model:

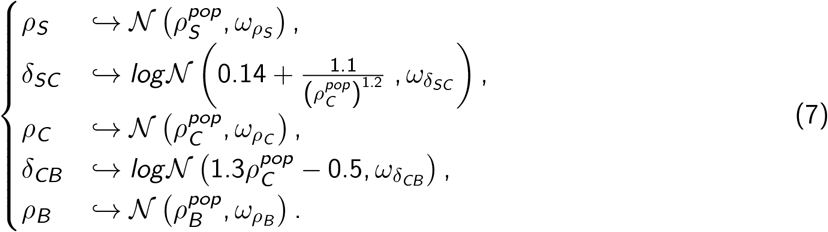

Following the same approach as for Model (2), we ran the SAEM algorithm on this model 50 times using uniformly sampled initial guesses for the population parameters. The resulting optimal likelihood distribution is displayed on Figure S1A. Most of the runs reached the same likelihood optimum as with Model (2) (Figure 3A), but 10 of them found higher likelihood values. In the case of Model (7), Akaike’s weights select only 36 runs as the best ones (Figure S1B), that we will consider as the runs that reached the global likelihood optimum.

The parameter values estimated in these 36 runs are displayed on Figure 4D. First, it shows that the reduction of the model did not affect the accuracy of the estimation for the five parameters that were identifiable in Model (2). Then, the population parameters 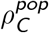 and 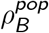 are estimated more precisely in the reduced model (7) than in Model (2). However the three standard deviations 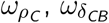 and 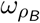 are still estimated with some variability.

Since 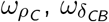 and 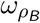 define the distributions of three random effects, their unidenfiability might indicate an overparameterization of the random effects. We investigate this using the *η*-shrinkage of the individual random effects in the next section.

#### 3.2.2 Random effects: shrinkage

We measured the *η*− shrinkage in the convergent runs of Model (7). The average shrinkage values for each parameter are displayed in Table 1, which confirms that the individual parameters of Model (7) are unidentifiable

**Table 1:**
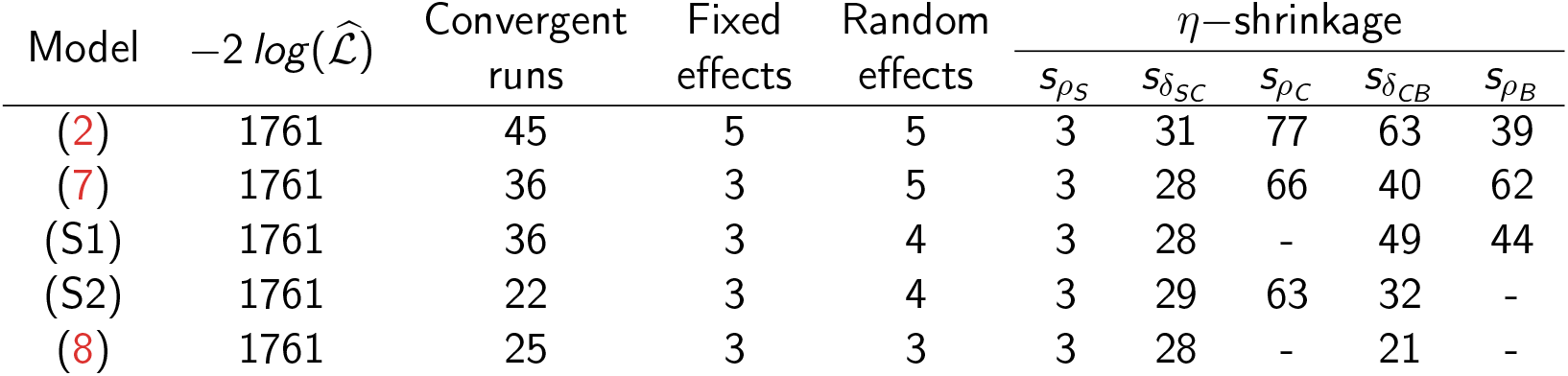
A comparison of our models along reduction. The table displays the optimal likelihood, the number of convergent runs (as selected by Akaike’s weights over 50 SAEM runs), the number of fixed and random effects, as well as the average *η*–shrinkage for each parameter (expressed as a percentage of the population variance) over the convergent runs for our models. For each model, we give the reference of the equation where it is defined.

These results indicate the individual data that we presented in Figure 1 are insufficient to estimate five parameters per individual precisely. Indeed, our dataset only comprises 7 individuals. In order to obtain an identifiable model based on Model (7), one might remove the random effect on one or several individual parameters. Fixing their values across the population might allow for a more precise estimation of the other, still variable, individual parameters, while keeping the same quantitative fit as with Models (2) and (7). However, all parameters are not necessarily equivalent in this regard, since different parameter sensitivities would make the model output more flexible under some combinations of fixed parameters. This would allow for these combinations to better fit the data, depending on the sensitivies of the model output to the parameter values. This sensitivity is imposed by the analytical solution to the structural equations of the model that we defined in System (1), but for most models there is no closed-form expression for the parameter sensitivities. As a consequence, and in order to keep our approach as general as possible, we will not attempt any analytical expression of the model output sensitivities to the individual parameters herein.

In order to choose which random effect to remove from our model, we consider that the parameter with the highest shrinkage is the most poorly estimated across the population. Since *ρ*_*C*_ and *ρ*_*B*_ display similar amounts of shrinkage in Model (7), we might remove either of their random effects in order to reduce our model. Removing the random effect on *ρ*_*C*_ defines a new model with reduced 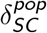 and 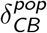, and with no variability on *ρC*, which is described in System (S1) in the Supplementary Materials. Conversely, removing the random effect on *ρ*_*B*_ defines a new model with reduced 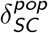 and 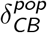, and with no variability on *ρ*_*B*_, which is described in System (S2) in the Supplementary Materials.

In both Model (S1) and Model (S2), the population parameters (Figures S4 and S7) and the individual parameters (Table 1, Figures S5 and S8) are still unidentifiable. We conclude that removing one random effect from Model (7) is not sufficient to make it identifiable. Thus, we propose to further reduce it by removing both the random effects on *ρ*_*C*_ and on *ρ*_*B*_, resulting in a reduced model with 3 remaining random effects:

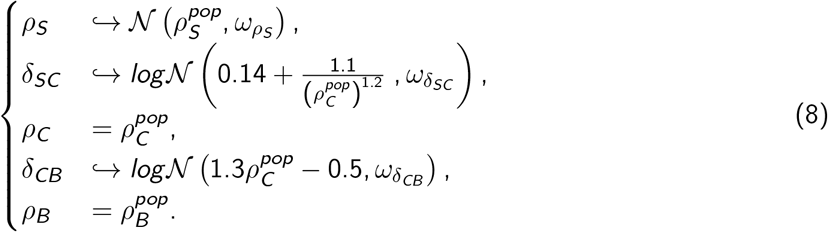

In this model, every population parameters –including the remaining variances 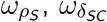 and 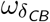– are reliably estimated (Figure S10) and the average shrinkage is lower than 30% for every parameter (Table 1, Figure S11). In other words, our reduction approach allowed us to define a fully identifiable MEM, *i*.*e*. a model which is able to quantitatively reproduce the individual trajectories of Figure 1, while explaining experimental heterogeneity in terms of precise parameter variations between individuals.

In the next section, we test another hypothesis, under which experimental heterogeneity does not come from inter-individual variations of the parameters of the dynamic model, but rather from variations in the initial condition of the experiment. Finally, we discuss the biological significance of the parameter values of Model 8 in Section 3.4.

### 3.3 Variability of the initial condition

In the previous section, we have considered that experimental heterogeneity originates from individual differences in the parameters of proliferation and differentiation kinetics. On the other hand, experimental heterogeneity might also be caused by an error in the sampling of the inital 25000 cells in the culture. In this section, we assess whether a variability of the initial condition could better account for experimental heterogeneity than variability on the model dynamic parameters. This can be tested by defining Mixed Effect Models accounting for the heterogeneity of the initial population size. First, we define three alternative versions of Model (8), which differ by their definition of the initial condition. We then calibrate these models and study their identifiability in order to select the best model at reproducing our data that is also identifiable.

The first model that we are considering is our reduced model (8), in which the kinetic parameters *ρ*_*S*_, *δ*_*SC*_ and *δ*_*CB*_ can vary between individuals, while *ρ*_*C*_ and *ρ*_*B*_ are kept constant between individuals. In this model, the initial condition is fixed for all individuals: (*S*_0_, *T*_0_, *B*_0_) = (25000, 25000, 0).

In the second model that we consider, all kinetic parameters are fixed to their population average, and we allow the initial condition to vary between individuals:

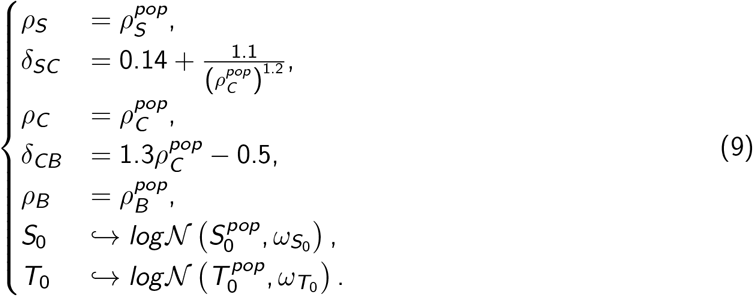

In this model, parameters 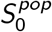 and 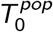 are the average of the initial number of cells in the LM1 and DM17 experiment. They represent a systematic error in the sampling of the initial 25000 cells. Parameters 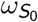 and 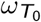 are the standard deviations of the initial number of cells in each experiment. For the third observable *B*, which is the number of differentiated cells, we consider the fixed initial condition *B*_0_ = 0 for all individuals, as the differentiation is initiated at time *t* = 0.

Finally, the last model that we consider allows for interindividual variations of both the kinetic parameters, as in Model (8), and the initial condition, as in Model (9):

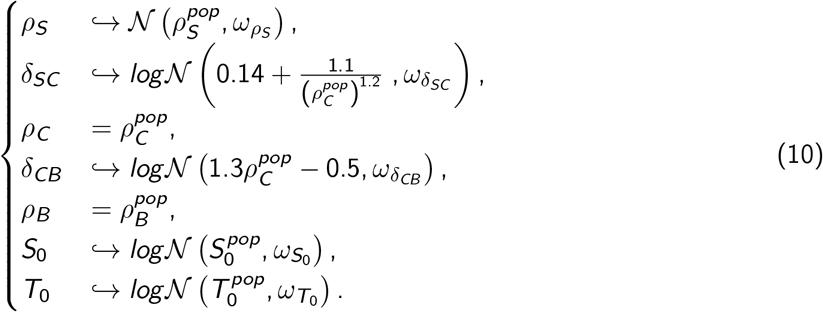

We present the convergence data and the distributions of the population parameters and shrinkage values for Models (9) and (10) in Sections S4.1 and S4.2 respectively of the Supplementary Materials.

We display the optimal log-likelihood 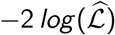 and the corresponding BIC for Models (8-10) in Table 2. Model (8) appears as the best one, closely followed by Model (10). On the other hand, Model (9) performs much worse than its competitors. Since Model (10) is unidentifiable (Figures S16 and S17), we conclude that Model (8) is the best one both in terms of quality of the fit and of parameter identifiability. This means that individual variations in the parameter values are more important in accounting for experimental heterogeneity than variations in the initial condition.

**Table 2:** Bayesian Information Criterion (BIC) computed for Models (8-10). Since the definition of the BIC depends on the decomposition of individual parameters between fixed p arameters a nd random parameters [37, 38], the computation of the BIC is ambiguous for Models (8) and (10). In these models, 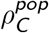 determines jointly the population values of *ρ*_*C*_ (which is fixed), *δ*_*SC*_ and *δ*_*CB*_ (which are random). Consequently, we can compute two different values of the BIC, depending on whether we consider 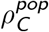 as the fixed effect of a fixed parameter (a), or as the fixed effect of two random parameters (b). In practice, this consideration does not seem to affect the outcome of the selection.

| Model | $2\ln(\hat{\mathcal{L}})$ | $\#\theta_F$ | $\#\theta_R$ | BIC |
| --- | --- | --- | --- | --- |
| (8) | 1761 | $2^a/1^b$ | $4^a/5^b$ | $1778^a/1775^b$ |
| (9) | 1801 | 3 | 4 | 1822 |
| (10) | 1759 | $2^a/1^b$ | $8^a/9^b$ | $1784^a/1782^b$ |

### 3.4 The final model

In Model (8), the population parameters are identifiable (Figure 5). It is the same for the individual parameters (Figure S11). This means that every parameter of the model can be reliably estimated from our data, and that the estimated values reflect an actual optimum in the description of *in vitro* erythropoiesis.

**Figure 5:**
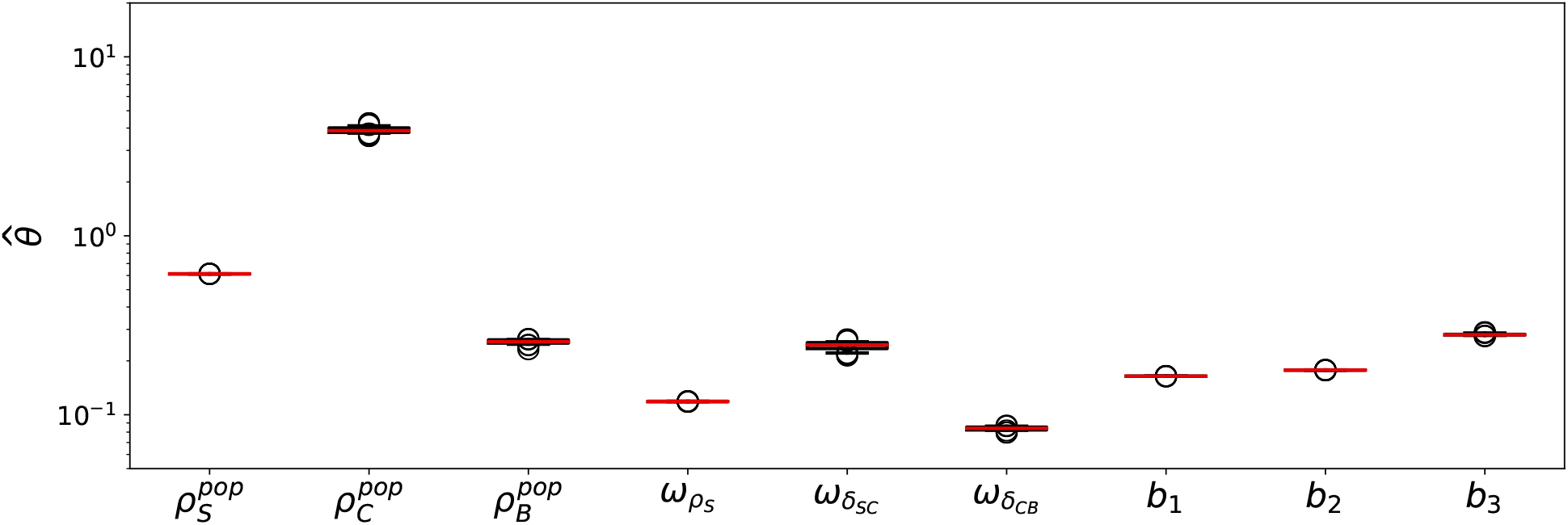
The estimated parameter values of Model (8) are identifiable. Displayed are the distributions of estimated population parameter values in the 25 convergent runs of SAEM for Model (8). The population means and standard deviations are expressed in *d*^−1^. The error parameters are dimensionless.

The population average of the parameter values are displayed in Table 3. The fixed effects 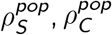, and 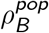 determine the average behavior of the experiment. The average proliferation rate 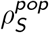 is estimated at 0.61 d^−1^ (Table 3). The doubling time of the self-renewing cells (*i*.*e*. the time it would take to double their population in the absence of differentiation) is thus 27 h in the average experiment, which is longer than the originally reported 18 h [33]. Proliferation in the committed compartment is much faster (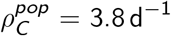, Table 3), which gives the committed cells an approximate doubling time of 4 h. Even though T2EC cells are known to proliferate faster in the differentiation medium than in the self-renewal medium [33], such a difference in proliferation times is rather intriguing.

**Table 3:** Parameter values in the optimal-likelihood run of Model (8). The table displays, for each parameter, the average value across the population in the SAEM run with the lowest 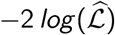. For the three parameters that vary across the population, we also give the bounds of the confidence intervals at level 0.95 for the individual parameter values. All values expressed in d^−1^

| Parameter | Lower bound | Population average | Upper bound |
| --- | --- | --- | --- |
| $\rho_S$ | 0.38 | 0.61 | 0.85 |
| $\delta_{SC}$ | 0.22 | 0.37 | 0.59 |
| $\rho_C$ | - | 3.8 | - |
| $\delta_{CB}$ | 3.8 | 4.5 | 5.3 |
| $\rho_B$ | - | 0.26 | - |

Moreover, 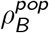 is estimated at 0.26 d^−1^, giving the differentiated cells a doubling time of 65 h. This means that their proliferation is almost invisible at the timescale of the experiments, as might be expected from differentiated cells.

From Equation (5), the value of 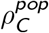 sets *δ*_*SC*_ to an average 0.37 d^−1^. The half-life of the self-renewing cells (*i*.*e*. the time it would take to differentiate half of the population in the absence of proliferation) is thus approximately 45 h. Respectively, from Equation (6), the average value of *δ*_*CB*_ in the population is estimated at 4.5 d^−1^, which gives the committed cells a half-life of approximately 3 h in the average experiment.

Apart from this average behaviour, three parameters of the final model can vary across the population, and are estimated at different values for each individual experiment. The first one is *ρ*_*S*_, which has the estimated variance 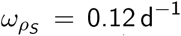. This translates into the individual values of *ρ*_*S*_ being estimated between 0.38 d^−1^ and 0.85 d^−1^ (Table 3), which corresponds to doubling times between 20 h and 44 h. Then, 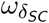 is estimated at 0.25 d^−^ with the individual parameter values of *δ*_*SC*_ estimated from 0.22 d^-1^ to 0.59 d^−1^, which means that the corresponding half-life ranges from 28 h to 76 h approximately. Finally, 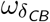 is estimated at 0.086 d^−1^, with individual parameter values for *δ*_*CB*_ ranging from 3.8 d^−1^ to 5.3 d^−1^ and the corresponding half-life approximately ranging from 3 h to 4h30.

## 4 Discussion and prospects

### 4.1 Generalizing our approach

Most approaches for identifiability analysis in MEM rely upon the FIM [10, 13, 14, 21–23], even though its parabolic approximation of the likelihood surface might mask complex practical unidentifiabilities [6]. We rather propose a multistart approach [13], that we refer to as *Initial Guess Sampling*. Multistart approaches do not provide any information regarding the confidence intervals of the model parameters [20], but they indicate parameter unidentifiabilities when the estimated values differ. For this reason, our samples of estimated values cannot be used in statistical analyses. On the contrary, they only allow for a visual check of the estimated values (for instance on Figure 5), thus adding a new kind of diagnostic plot to the visual tools already available to the modeller using mixed effect models.

This approach allowed us to design an identifiable MEM of *in vitro* erythropoiesis which accounts well for experimental heterogeneity as inter-individual variations of the proliferation and differentiation parameters. Then, a question that naturally arises is whether or not our approach could be applied, or generalized, onto other MEM? We identified two features of our approach, that might be of importance for such a generalization.

First, it seems that iteratively reducing our model affects the convergence of SAEM. Indeed, while all 50 runs of SAEM reached the same likelihood optimum for Model (2) (Figure 3A), only 36 runs reached it for Model (7) (Figure S1B). Convergence was further affected by removing random effects from our model (Figures S3B & S6B), to the point where only 25 runs of SAEM reached the likelihood optimum for Model (8) (Figure S9B). However, the number of convergent SAEM runs is critical to the assessment of population parameters identifiability. Depending on the complexity of the model and the dataset, 50 SAEM runs might not be sufficient to assess parameter identifiability and allow for model reduction, and the number of runs to be performed should thus be finely tuned in order to avoid issues with computational time.

Morevover, we used the correlations between population parameters to define Model (7), with constraints on the fixed effects. The exact shape of the likelihood landscape and the resulting unidentifiability is related to the structure of the model, and the quality of the data. This means that we were able to explore the parameter space near the likelihood optimum using pairwise correlations (Figure 4). Yet, in more complex nonlinear MEM, it is possible that the correlations would involve more than two parameters at a time. In the end, detecting these complex correlations would require some kind of multivariate correlation analysis [41].

### 4.2 About the source of experimental heterogeneity

In this paper, we consider that experimental heterogeneity might either originate in variations of the kinetic parameters between replicates of the experiment, or by experimental errors in the initial number of cells. Using model selection and identifiability analysis, we conclude that variations in the kinetic parameters of proliferation and differentiation best explain experimental heterogeneity.

Considering that every replicate of the experiment was obtained with the exact same protocol (Section 2), it seems that only two features of our experiment could change from replicate to replicate.

The first one is the group of 25000 cells used to initiate the culture. In the haematopoietic system, *in vivo* stem cells and progenitors display substancial variations in terms of self-renewal and potency [42]. Since our T2EC cells are erythropoietic progenitors, our results suggest that the self-renewal and differentiation abilitites vary between the cell populations that we used to initiate every experiment.

On the other hand, there has also been discussion around the fact that the external temperature of the incubators (*i*.*e*. the temperature of the room where the T2EC are incubated, which is not their incubation temperature) might affect the variability of gene expression [43]. This in turn, could affect their self-renewal or differentiation potency.

### 4.3 Conclusion

In this paper, we proposed a MEM for *in vitro* erythropoiesis, that accounts for experimental heterogeneity. We developped a multistart approach for assessing its identifiability, and we successfully reduced it to make it identifiable. We showed that experimental heterogeneity is faithfully accounted for by variations of the kinetic parameters of proliferation and differentiation in our system, and we relate these parameter variations to actual biological features of our cells. This work establishes a MEM framework to study variability in the outcome of biological experiments. Furthermore, it proposes a novel approach for the analysis of parameter identifiability in MEM, and for reducing unidentifiable MEM.

## Declarations

## Acknowledgements

We thank the members of the Systems Biology of Decision Making team at the Laboratory of Biology and Modelling of the Cell, as well as the Dracula team of Inria, for insightful discussion all along the course of this project.

We thank the BioSyL Federation and the LabEx Ecofect (ANR-11-LABX-0048) of the University of Lyon for inspiring scientific events.

We also would like to thank members of the Population Approach Group Europe that were present at the PAGE meeting 2019 for enlightening feedback.

Finally, we want to thank Pr. Saccomani (University of Padova, Italy) for providing the manuscript for reference [9].

## Availability of data and materials

All the datasets and pieces of code analysed and generated during the current study are available in a public github repository, at https://github.com/rduchesn/MixedEffectModelReduction.

## Ethics approval and consent to participate

All methods were carried out in accordance with relevant guidelines and regulations, notably the DIRECTIVE 2010/63/EU OF THE EUROPEAN PARLIAMENT AND OF THE COUNCIL of 22 September

2010 regarding “the killing of animals solely for the use of their organs or tissues.” (https://eur-lex.europa.eu/legal-content/EN/TXT/HTML/?uri=CELEX:32010L0063)

## Authors contributions

AG generated the experimental data used in this study. RD designed the models, estimated their parameter values, carried out all subsequent analyses, and wrote the first draft of the manuscript. FC and OG supervised the project. All authors read and reviewed the manuscript, and they approved its final version.

