## Supplementary Materials for "Practical Identifiability in the Frame of Nonlinear Mixed Effects Models: the Example of the *in vitro* Erythropoiesis"

All the datasets and pieces of code analysed and generated during the current study are available in a public github repository, at https://github.com/rduchesn/MixedEffectModelReduction.

#### S1 Composition of the culture media

| LM1 | | DM17 | |
|---|---|---|---|
| Component | Concentration | Component | Concentration |
| $\alpha$-MEM medium | - | $\alpha$-MEM medium | - |
| Foetal bovine serum (FBS) | 10 % | Foetal bovine serum (FBS) | 10 % |
| HEPES | 1 mM | HEPES | 1 mM |
| $\beta$-mercaptoethanol | 100 nM | $\beta$-mercaptoethanol | 100 nM |
| penicillin | 100 U/mL | penicillin | 100 U/mL |
| streptomycin | 100 U/mL | streptomycin | 100 U/mL |
| TGF-$\alpha$ | 5 ng/mL | insulin | 10 ng/mL |
| TGF-$\beta$ | 1 ng/mL | anaemic chicken serum (ACS) | 5% |
| dexamethasone | 1 mM | | |

Table S1: Composition of the culture media. Both the LM1 and the DM17 media are obtained by supplementing an $\alpha$-MEM medium with various chemicals that allow for the proliferation (LM1) and differentiation (DM17) of T2EC cells.

#### S2 Parameter estimation

| Task | Parameter | Value |
|---|---|---|
| Population parameters | Number of iterations (burnin phase) | 5 |
| | Number of iterations (exploratory phase) | 5000[a] |
| | Simulated annealing decrease rates | 0.95 |
| | Number of iterations (smoothing phase) | 1000[a] |
| | Stepsize exponent | 0.7 |
| Individual parameters (Conditional Mean) | Interval length | 50 |
| | Relative interval width | 0.05 |
| | Simulated parameters per individual | 10 |
| FIM estimation (Stochastic Approximation) | Minimum number of iterations | 50 |
| | Maximum number of iterations | 200 |
| Likelihood estimation (Importance Sampling) | Monte Carlo size | 10 000 |
| | Degrees of freedom of the t distribution | $\{1, 2, 5, 10, 15\}$ |

Table S2: SAEM parameters. When Monolix implements several different methods to perform a task, we also indicate which method we used between brackets. a: Non-default parameter values

| Parameter | Initial guess | Unit |
|---|---|---|
| $\rho_S^{pop}$ | $\mathcal{U}(-5, 5)$ | $d^{-1}$ |
| $\delta_{SC}^{pop}$ | $\mathcal{U}(0, 5)$ | $d^{-1}$ |
| $\rho_C^{pop}$ | $\mathcal{U}(-5, 5)$[a] | $d^{-1}$ |
| $\delta_{CB}^{pop}$ | $\mathcal{U}(0, 5)$ | $d^{-1}$ |
| $\rho_B^{pop}$ | $\mathcal{U}(-5, 5)$ | $d^{-1}$ |
| $\omega_{\rho_S}$ | 5 | $d^{-1}$ |
| $\omega_{\delta_{SC}}$ | 5 | $d^{-1}$ |
| $\omega_{\rho_C}$ | 5 | $d^{-1}$ |
| $\omega_{\delta_{CB}}$ | 5 | $d^{-1}$ |
| $\omega_{\rho_B}$ | 5 | $d^{-1}$ |
| $b_1$ | 1 | - |
| $b_2$ | 1 | - |
| $b_3$ | 1 | - |

Table S3: Initial guess values and distributions in our models. The initial guesses of the fixed effects are sampled uniformly with respect to each parameter bounds. The initial guess of the variances of the random effects, and of the error parameters, are set to an arbitrary high value to improve the convergence of SAEM in the early phase of the estimation. a: In the models where $\delta_{SC}^{pop}$ and $\delta_{CB}^{pop}$ are defined with respect to the value of $\rho_C^{pop}$ by equations (5) and (6), the initial guess of $\rho_C^{pop}$ is sampled uniformly in $[0.39, 5]$ to ensure the positivity of $\delta_{SC}^{pop}$ and $\delta_{CB}^{pop}$.

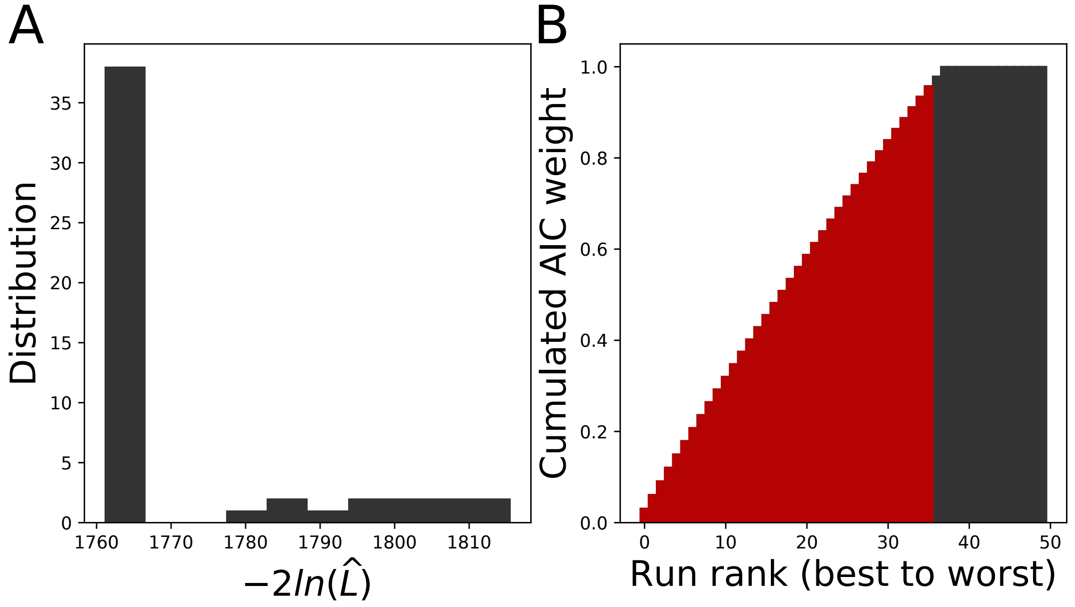


Figure S1: Likelihood distribution over 50 SAEM runs on Model (7) (A) and the corresponding cumulated AIC weights (B). The 36 runs associated to the lowest likelihood values (*i.e.* those that add up to 95% of the total weight of the 50 runs) are coloured in red.

### S3 Reduced Models

#### S3.1 Model with reduced $\delta_{SC}^{pop}$ and $\delta_{CB}^{pop}$

The model with reduced $\delta_{SC}^{pop}$ and $\delta_{CB}^{pop}$ is defined in System (7) in the main manuscript. Over the 50 runs of SAEM that we performed, Akaike's weights select 36 as the best-ones. We consider them as those that reached the global likelihood optimum. As discussed in the main text, its population parameters are unidentifiable (Figure 4D). The values of the $\eta$-shrinkage over the 36 convergent are displayed on Figure S2. They confirm that Model (7) is also unidentifiable at the individual scale.

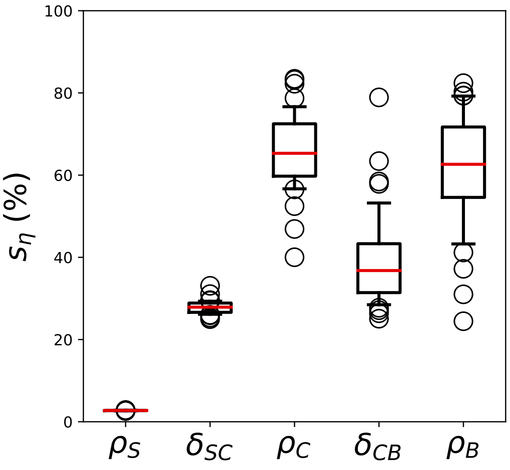


Figure S2: Distribution of $s_\eta$ values for the individual parameters in the 36 convergent runs of SAEM for Model (7).

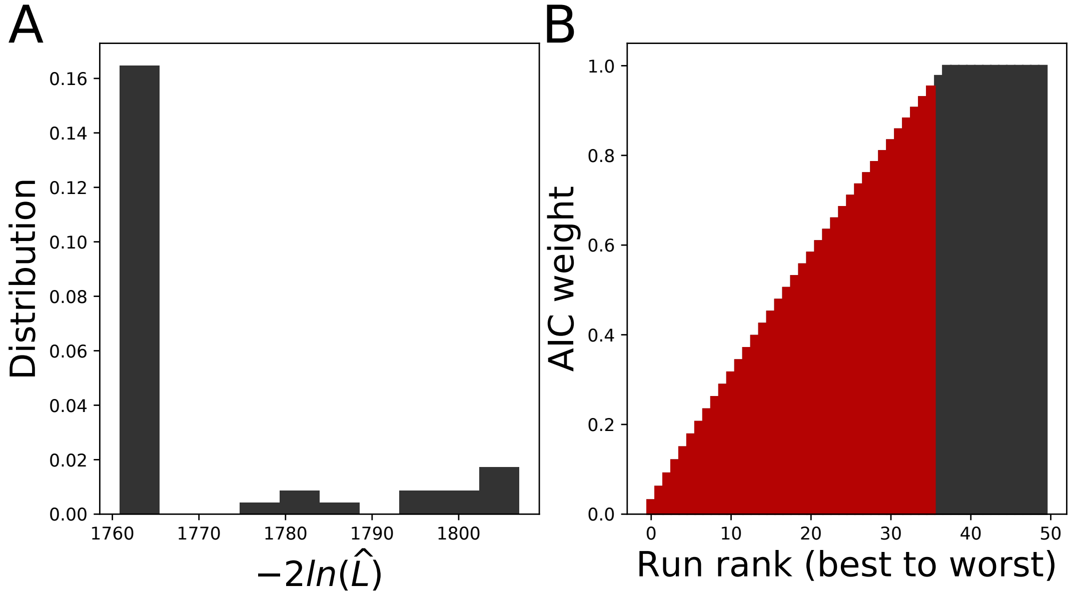


Figure S3: Likelihood distribution over 50 SAEM runs on Model (S1) (A) and the corresponding cumulated AIC weights (B). The 36 runs associated to the lowest likelihood values (*i.e.* those that add up to 95% of the total weight of the runs) are coloured in red.

##### S3.2 Model with reduced $\delta_{SC}^{pop}$, $\delta_{CB}^{pop}$ and a fixed $\rho_C$

Removing the individual effect on $\rho_C$ in our previous model leads to a model with one constant parameter over the population:

$$\begin{cases} \rho_S & \hookrightarrow \mathcal{N}\left(\rho_S^{pop}, \omega_{\rho_S}\right), \\ \delta_{SC} & \hookrightarrow log\mathcal{N}\left(0.14 + \frac{1.1}{\left(\rho_C^{pop}\right)^{1.2}}, \omega_{\delta_{SC}}\right), \\ \rho_C & = \rho_C^{pop}, \\ \delta_{CB} & \hookrightarrow log\mathcal{N}\left(1.3\rho_C^{pop} - 0.5, \omega_{\delta_{CB}}\right), \\ \rho_B & \hookrightarrow \mathcal{N}\left(\rho_B^{pop}, \omega_{\rho_B}\right). \end{cases} \tag{S1}$$

The distribution of the optimal likelihoods over 50 SAEM runs are displayed on Figure S3A. Akaike's weights select 36 runs as the best ones(Figure S3B), which we consider as those that reached the global likelihood optimum. For these 36 runs, the distribution of estimated population parameters is displayed on Figure S4). It shows that the population variances $\omega_{\delta_{CB}}$ and $\omega_{\rho_B}$ are unidentifiable, which might indicate an overparameterization of the random effects. Figure S5 displays the shrinkage distributions over the convergent runs, confirming that the individual parameters $\delta_{CB}$ and $\rho_B$ are unidentifiable.

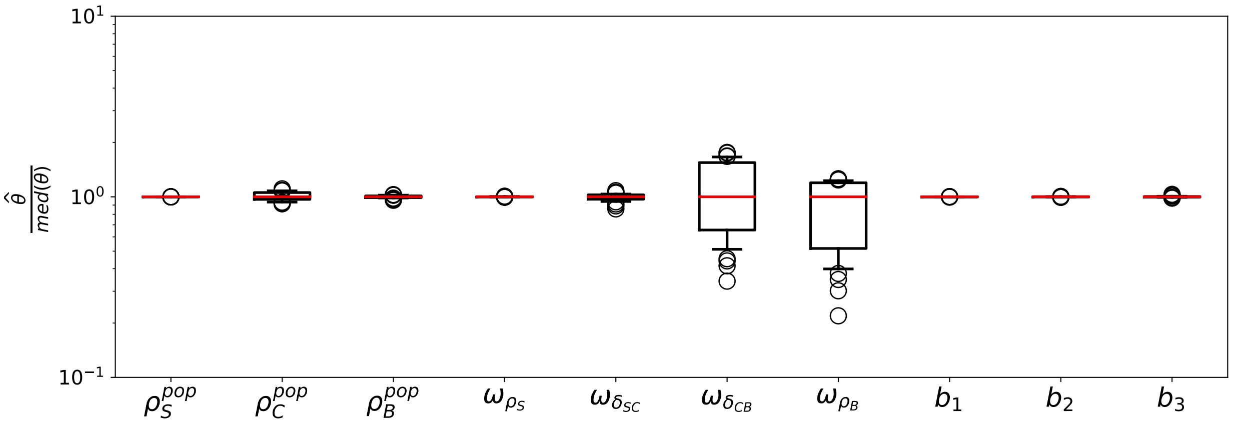


Figure S4: Estimated parameter values in the 36 convergent runs of SAEM for Model (S1). Displayed are the distributions of estimated parameter values, normalized by their median.

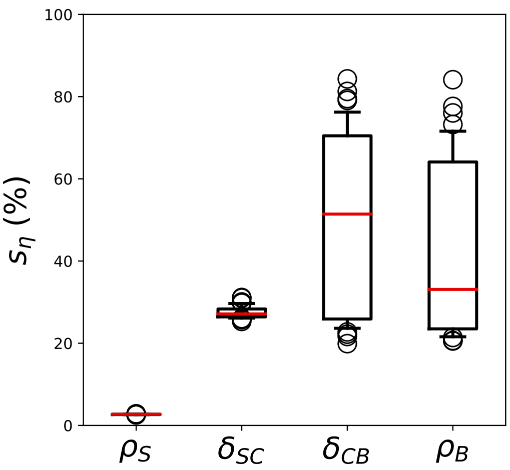


Figure S5: Distribution of $s_\eta$ values for the individual parameters in the 36 convergent runs of SAEM of Model (S1).

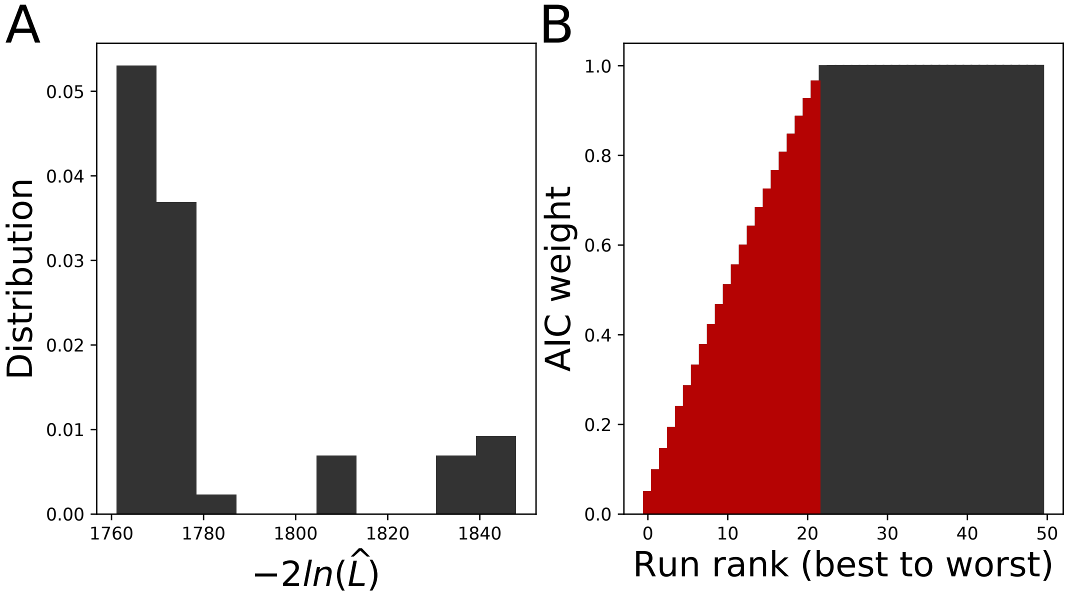


Figure S6: Likelihood distribution over 50 SAEM runs on Model (S2) (A) and the corresponding cumulated AIC weights (B). The 22 runs associated to the lowest likelihood values (*i.e.* those that add up to 95% of the total weight of the runs) are coloured in red.

#### S3.3 Model with reduced $\delta_{SC}^{pop}$, $\delta_{CB}^{pop}$ and a fixed $\rho_B$

Removing the individual effect on $\rho_B$ in Model (7) leads to a model with one constant parameter over the population:

$$\begin{cases} \rho_S & \hookrightarrow \mathcal{N}\left(\rho_S^{pop}, \omega_{\rho_S}\right), \\ \delta_{SC} & \hookrightarrow log\mathcal{N}\left(0.14 + \frac{1.1}{\left(\rho_C^{pop}\right)^{1.2}}, \omega_{\delta_{SC}}\right), \\ \rho_C & =\hookrightarrow \mathcal{N}\left(\rho_C^{pop}, \omega_{\rho_C}\right), \\ \delta_{CB} & \hookrightarrow log\mathcal{N}\left(1.3\rho_C^{pop} - 0.5, \omega_{\delta_{CB}}\right), \\ \rho_B & = \rho_B^{pop}. \end{cases} \tag{S2}$$

The distribution of the optimal likelihoods over 50 SAEM runs are displayed on Figure S6A. Akaike's weights select 22 runs as the best ones(Figure S6B), which we consider as those that reached the global likelihood optimum. For these 22 runs, the distribution of estimated population parameters is displayed on Figure S7). It shows that the population variances $\omega_{\rho_C}$ and $\omega_{\delta_{CB}}$ are unidentifiable, which might indicate an overparameterization of the random effects. Figure S8 displays the shrinkage distributions over the convergent runs, confirming that the individual parameters $\rho_C$ and $\delta_{CB}$ are unidentifiable, as well as $\delta_{SC}$.

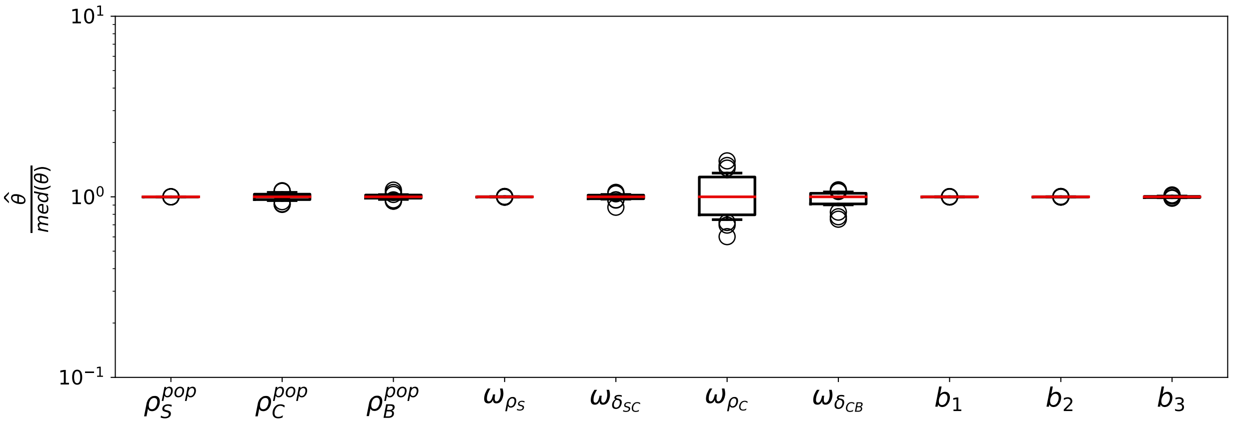


Figure S7: Estimated parameter values in the 22 convergent runs of SAEM for Model (S2). Displayed are the distributions of estimated parameter values, normalized by their median.

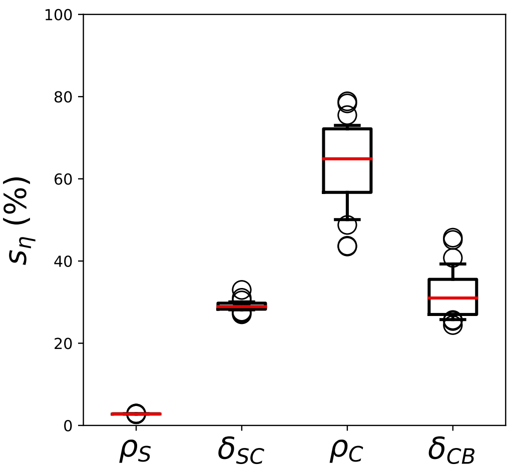


Figure S8: Distribution of $s_\eta$ values for the individual parameters in the 22 convergent runs of SAEM of Model (S2).

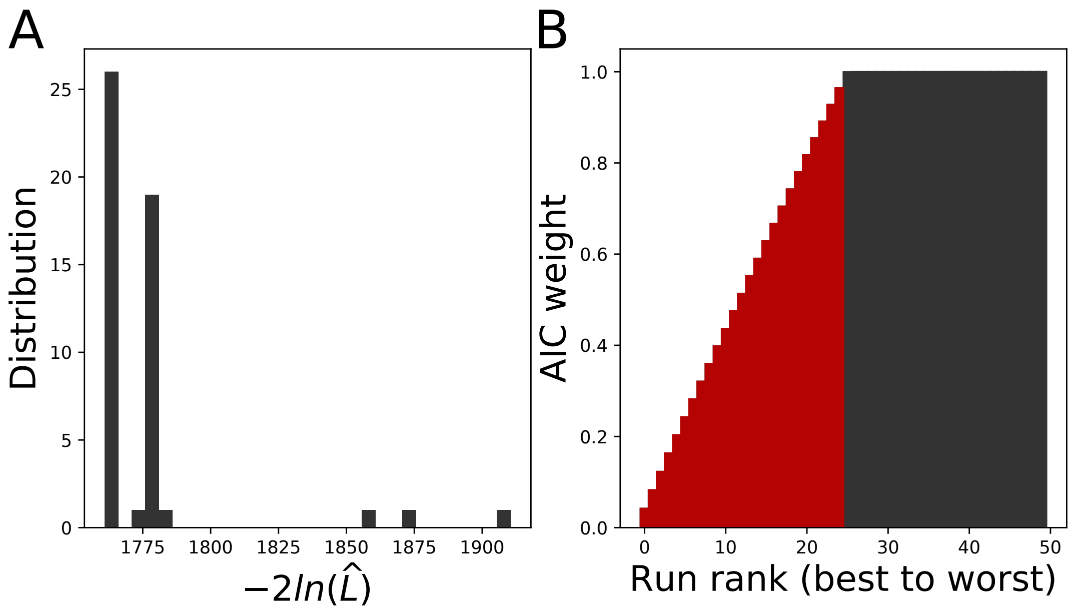


Figure S9: Likelihood distribution over 50 SAEM runs on Model (8). (A) and the corresponding cumulated AIC weights (B). The 25 runs associated to the lowest likelihood values (*i.e.* those that add up to 95% of the total weight of the 50 runs) are coloured in red.

##### S3.4 Model with reduced $\delta_{SC}^{pop}$, $\delta_{CB}^{pop}$ and fixed $\rho_C$ and $\rho_B$

We define Model (8) by removing the random effects on $\rho_C$ and $\rho_B$ from Model (7). We display the distribution of the estimated likelihood values over 50 SAEM runs on Figure S9A. The global likelihood minimum is located, as with the previous models, around $-2\,log(\widehat{\mathcal{L}}) = 1761$. Interestingly, there is a second peak in the likelihood distribution, located around $-2\,log(\widehat{\mathcal{L}}) = 1777$. Yet Akaike's weights only select the first peak as the global likelihood optimum, as they select 25 runs among the 50 (Figure S9B). Among these runs, the population parameters are reliably estimated (Figure S10), and the shrinkage levels indicate an overall good agreement of the individual and population parameter distributions (Figure S11).

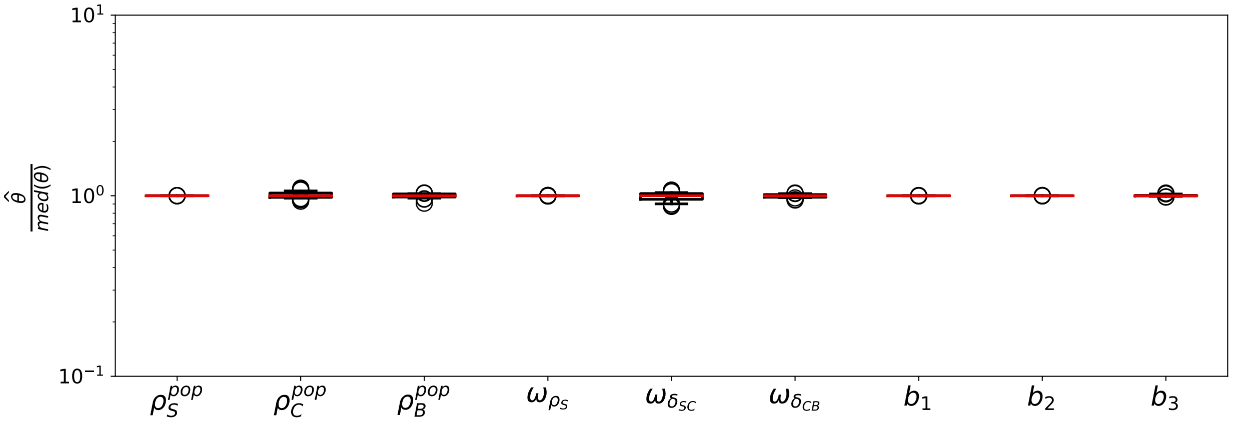


Figure S10: Estimated parameter values in the 25 convergent runs of SAEM for Model (8). Displayed are the distributions of estimated parameter values, normalized by their median.

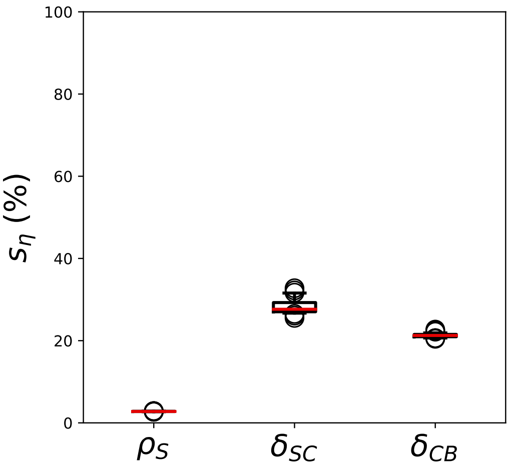


Figure S11: Distribution of $s_\eta$ values for the individual parameters in the 25 convergent runs of SAEM for Model (8).

### S4 Variability on the initial condition

#### S4.1 Model with fixed kinetic parameters and variability on the initial condition

We define Model (9) by fixing the values of all parameters of the dynamic model across the population, and by allowing for a variation of the initial condition ($S_0$, $T_0$, $B_0$) between individuals. Akaike's weights select 24 of the 50 SAEM runs as the convergent ones (Figure S12), for which both the population parameters (Figure S13) and the individual parameters (Figure S14) appear as identifiable.

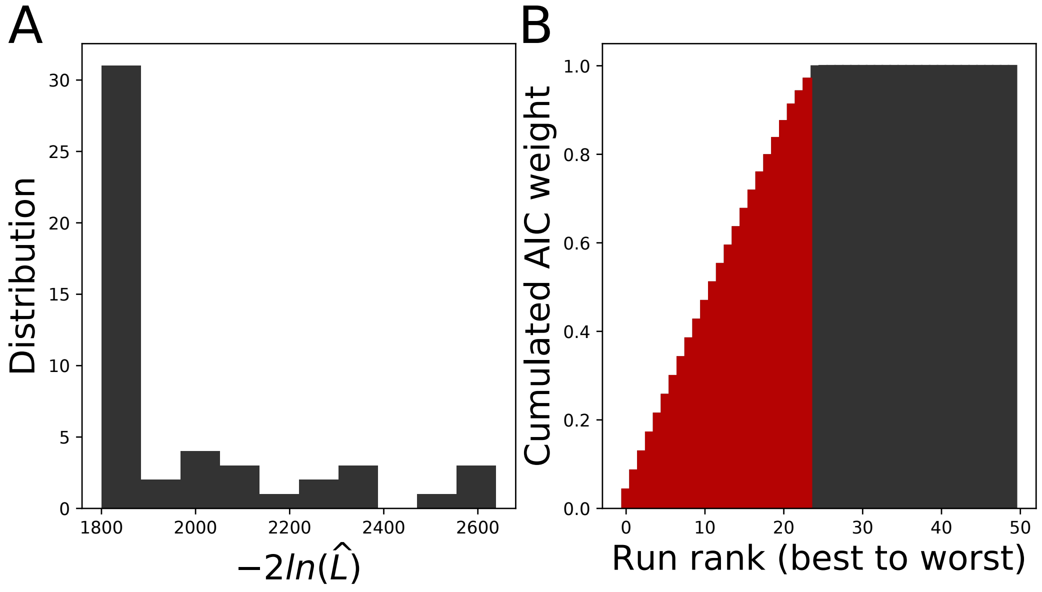


Figure S12: Likelihood distribution over 50 SAEM runs of Model (9). (A) and the corresponding cumulated AIC weights (B). The 24 runs associated to the lowest likelihood values (*i.e.* those that add up to 95% of the total weight of the 50 runs) are coloured in red.

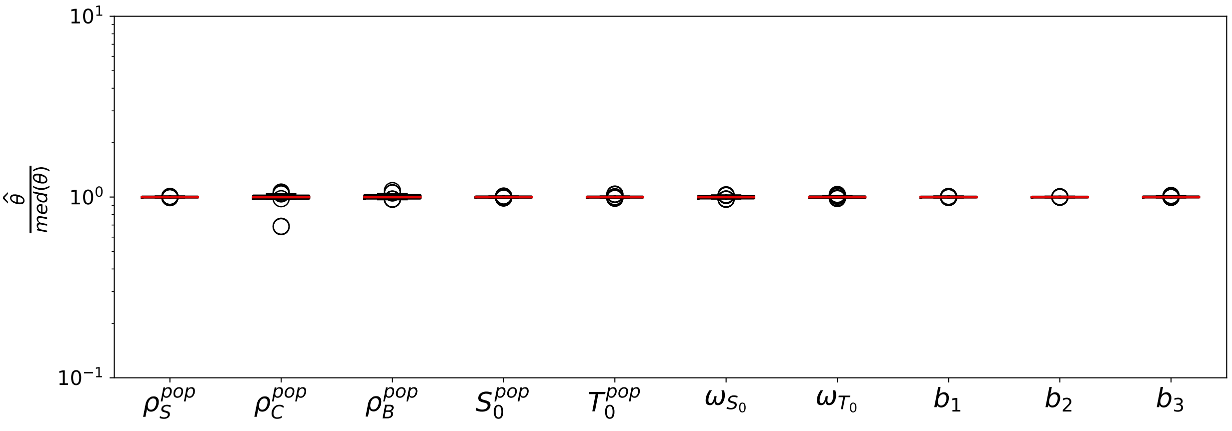


Figure S13: Estimated parameter values in the 24 convergent runs of SAEM of Model (9). Displayed are the distributions of estimated parameter values, normalized by their median.

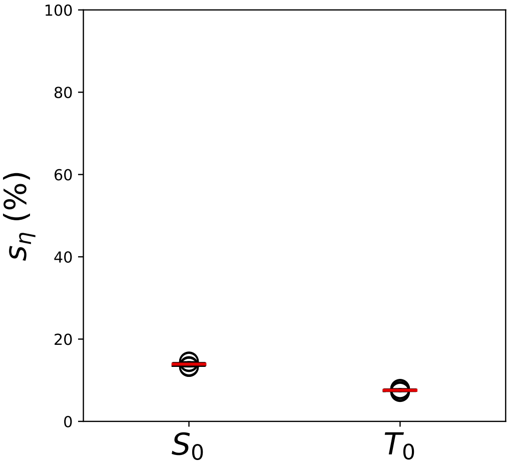


Figure S14: Distribution of $s_\eta$ values for the individual parameters in the 24 convergent runs of SAEM of Model (9)

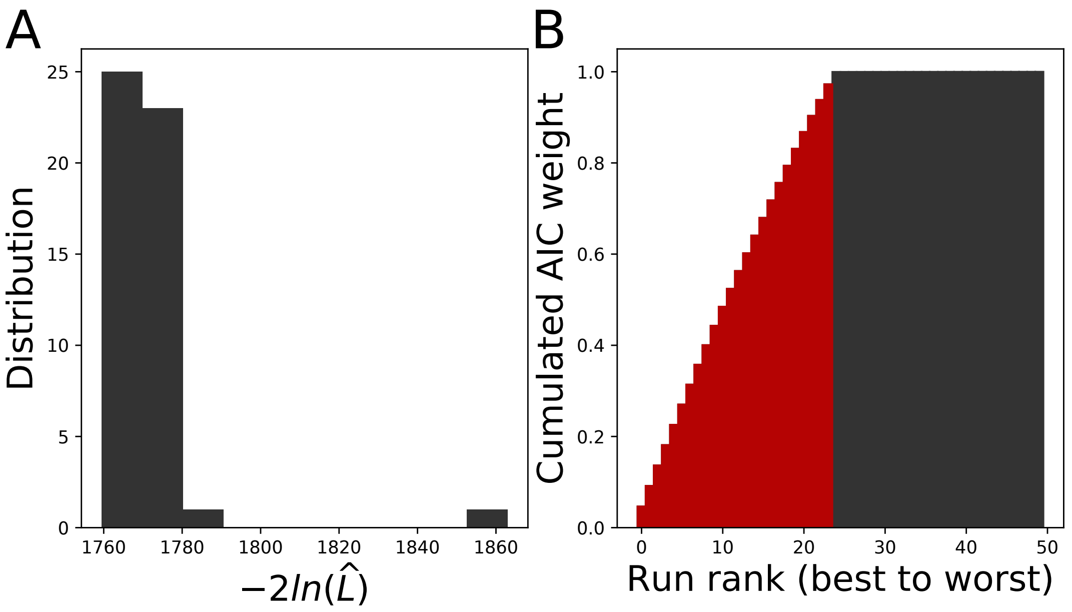


Figure S15: Likelihood distribution over 50 SAEM runs of Model (10). (A) and the corresponding cumulated AIC weights (B). The 24 runs associated to the lowest likelihood values (*i.e.* those that add up to 95% of the total weight of the 50 runs) are coloured in red.

##### S4.2 Model with variability on the kinetic parameters and the initial condition

We define Model (10) by allowing for a variation of the initial condition ($S_0$, $T_0$, $B_0$) between individuals in Model (8). Akaike's weights select 24 of the 50 SAEM runs as the convergent ones (Figure S15). In these runs, it appears that some population parameters, namely $\omega_{S_0}$ and $\omega_{T_0}$ (Figure S16), as well as the corresponding individual parameter (Figure S17) are unidentifiable. As a consequence, it appears impossible to infer at the same time the kinetic parameters and the initial condition of the model for all individuals. Moreover this model does not result in a improved BIC compared to Model (8), where the kinetic parameters can vary between individuals but not the initial condition (Table 2).

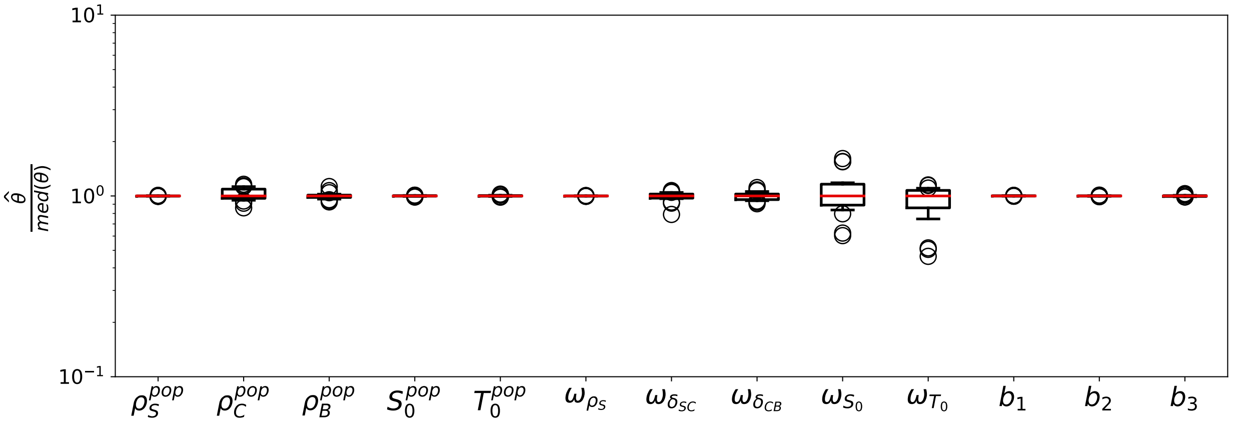


Figure S16: Estimated parameter values in the 24 convergent runs of SAEM of Model (10). Displayed are the distributions of estimated parameter values, normalized by their median.

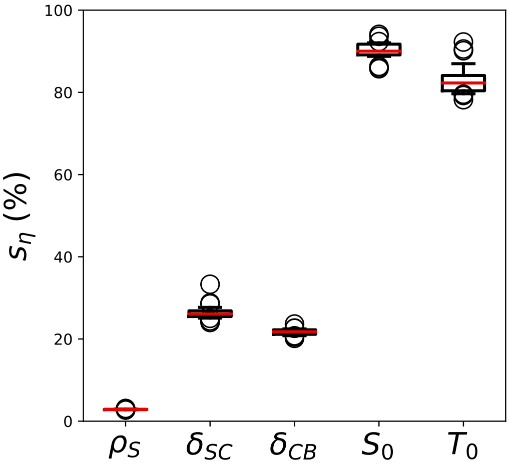


Figure S17: Distribution of $s_\eta$ values for the individual parameters in the 24 convergent runs of SAEM of Model (10).
